# *Escherichia coli* recombinant expression of SARS-CoV-2 protein fragments

**DOI:** 10.1101/2021.06.22.449540

**Authors:** Bailey E. McGuire, Julia E. Mela, Vanessa C. Thompson, Logan R Cucksey, Claire E. Stevens, Ralph L. McWhinnie, Dirk F. H. Winkler, Steven Pelech, Francis E. Nano

## Abstract

We have developed a method for the inexpensive, high-level expression of antigenic protein fragments of SARS-CoV-2 proteins in *Escherichia coli*. Our approach used the thermophilic family 9 carbohydrate-binding module (CBM9) as an N-terminal carrier protein and affinity tag. The CBM9 module was joined to SARS-CoV-2 protein fragments via a flexible proline-threonine linker, which proved to be resistant to *E. coli* proteases. Two CBM9-spike protein fragment fusion proteins and one CBM9-nucleocapsid fragment fusion protein largely resisted protease degradation, while most of the CBM9 fusion proteins were degraded at some site in the SARS-CoV-2 protein fragment. All fusion proteins were expressed in *E. coli* at about 0.1 g/L, and could be purified with a single affinity binding step using inexpensive cellulose powder. Three purified CBM9-SARS-CoV-2 fusion proteins were tested and found to bind antibody directed to the appropriate SARS-CoV-2 antigenic region. The largest intact CBM9 fusion protein incorporates spike protein amino acids 540-588, which is a conserved region immediately C-terminal to the receptor binding domain that is widely recognized by human convalescent sera and contains a putative protective epitope.

## INTRODUCTION

One of the public health surveillance tools needed to respond to the coronavirus disease 2019 (COVID-19) pandemic is the ability to detect seroconversion to antigens of the SARS-CoV-2 virus, the causative agent of COVID-19. The ability to detect antibodies that are specific to SARS-CoV-2 allows an assessment of the level of probable immunity to COVID-19 in a population. At the individual level, the ability to detect anti-SARS-CoV-2 antibodies can help one assess their personal level of vulnerability to COVID-19 due to immunity generated by either vaccination or from an indeterminate or asymptomatic infection.

The human antibody response to SARS-CoV-2 infection can include the development of antibodies reactive with any of the 29 proteins encoded in the viral genome (1), including 16 non-structural proteins (NSP’s) encoded by the ORF1a/b gene. However most studies have focused on studying the antibody response to the abundant spike and nucleocapsid proteins (2–11). Among the SARS-CoV-2 proteins, the spike protein varies the most between coronaviruses (12); using it allows for the greatest specificity in an antibody assay as well as the ability to differentiate reaction with SARS-CoV-2 from other common coronaviruses. As well some antibodies reactive with the spike protein, especially those that react within or near its receptor binding domain (RBD), are neutralizing (9, 11, 13–15). Thus, detection of anti-spike protein antibodies may indicate a level of immunity to COVID-19.

Some studies of anti-SARS-CoV-2 antibody response examine antibody reactivity with linear epitopes using synthetic peptides that correspond to the primary structure of viral proteins (2–10). While this type of assay misses many antibody responses against conformational and topographically assembled epitopes, it is the most practical, both technically and economically. Although synthetic peptides are far less expensive than full-length recombinant SARS-CoV-2 spike protein, their production cost can still present barriers in resource-poor health systems or when large quantities are needed.

The SARS-CoV-2 spike protein is the most common antigen used to test for seroconversion, and recombinant spike protein, the RBD fragment or synthetic peptides corresponding to spike sequences are commonly used to test sera. Typically, human derived HEK293 cells are used to express spike protein, and Stadlbauer et al. (16) reported expression levels of about 20 mg/L for the RBD fragment and about 5 mg/L of the trimer form of the full-length spike protein. However, other recombinant expression systems have been used to produce the SARS-CoV-2 spike protein. For example, Yang et al. (13) expressed the spike protein RBD in the Sf9 insect cell line (*Spodoptera frugiperda)* using a BAC-to-BAC expression system, and Fujita et al. (17) expressed full-length spike protein in silk worm larvae at a level of about 10 mg/L of larval serum. Recently Rihn et al. (18) described the construction of glutathione S-transferase (GST) and maltose binding protein (MBP) fusions to all of the ORFs of SARS-CoV-2, as part of an expansive effort to develop molecular tools to study SARS-CoV-2. These fusion proteins are expressed in *E. coli*, and while their properties and yields are not reported, these recombinants represent significant tools for obtaining SARS-CoV-2 antigenic material. In a similar fashion, our work explores the utility of expressing SARS-CoV-2 epitopes in *E. coli*, using the under-appreciated protein carrier family 9 carbohydrate-binding module (CBM9) from the *Thermotoga maritima* enzyme xylanase 10A (19, 20) that promotes high level protein expression and uses inexpensive reagents for protein purification (21, 22).

## MATERIALS AND METHODS

### Recombinant techniques

Plasmid pRSET5A was used as the backbone for all expression plasmid constructs (23). All of the synthetic DNA regions designed to encode CBM9-SARS-CoV-2 spike protein fusions were made by Twist Biosciences. To initially test the expression of CBM9 peptide fusions we cloned synthetic DNA encoding CBM9, CBM9-ID-C, CBM9-ID-F and CBM9-H3 (Fig. 1A). Plasmid pRSET5A was amplified by inverse PCR using primers F-R5A and R-R5A, which have Esp3I sites added to the ends that upon digestion yield 5’-overhangs compatible with the overhangs generated for the PCR amplicons of the synthetic DNA fragments. The CBM9 DNA fragment was designed with the native codon composition, since this was known to function in *E. coli* from previous work (20) and this fragment was amplified with primers F-nCBD and R-nCBD which have BsaI sites on their ends. Since the native *CBM9* gene has an internal Esp3I site and pRSET5A has internal BsaI sites, the plasmid and CBM9 amplicons were separately digested with Esp3I and BsaI and then joined by a standard DNA ligation. The CBM9-C, F and H3 DNA fragments were codon optimized (24) for *E. coli* and designed to lack an internal Esp3I site. These fragments were amplified with primers F-CBD (forward primer for all fragments) and R-CBD-IDc, R-CBD-IDf and R-CBD-h3 as the reverse primers. After amplification the products were joined to pRSET5A using a simultaneous cutting and ligation reaction (25) using Esp3I as the restriction enzyme. Briefly, 30 cycles of 5 min at 37 °C and 5 min at 16 °C were followed by 10 min at 65 °C. Ligated DNA was transformed into T7 Express *lysY/I^q^ E. coli* (NEB) and selected on LB agar (per liter, 5 g yeast extract; 10 g tryptone, 5 g NaCl; 15 g agar) supplemented with chloramphenicol (10 μg/mL) and carbenicillin (250 μg/mL). Once initial clones were sequence verified and shown to produce the appropriate protein product, further recombinants were constructed using the pRSET5A::*CBM9-id-c* clone as the backbone. This plasmid was amplified by inverse PCR using primers Fb-R5A and R-R5AidC so as to remove the SARS-CoV-2 spike protein-encoding fragment of DNA and replace it with another fragment of synthetic DNA (ID-a, b, d, g, h, h1, h2, i; Twist Biosciences) (Fig. 1B) using a cutting-ligation reaction as described above. To make a plasmid encoding just the CBM9-(TP)_4_P (no SARS-CoV-2 fragment) or CBM9-N (containing a nucleocapsid epitope), plasmid pRSET5A::*CBM9-id-a* was amplified with primers nF2-R5A-CBD and nR-R5A-Flex; or F-nucl-ep and R-nucl-ep and Esp3I digested and ligated with a strategy depicted in Fig. 1C. The primers used to make all of the constructs are listed in Supplemental Table S1. A color-coded example of a CBM9 fusion clone is shown in Supplemental Fig. S1.

**Figure 1.**
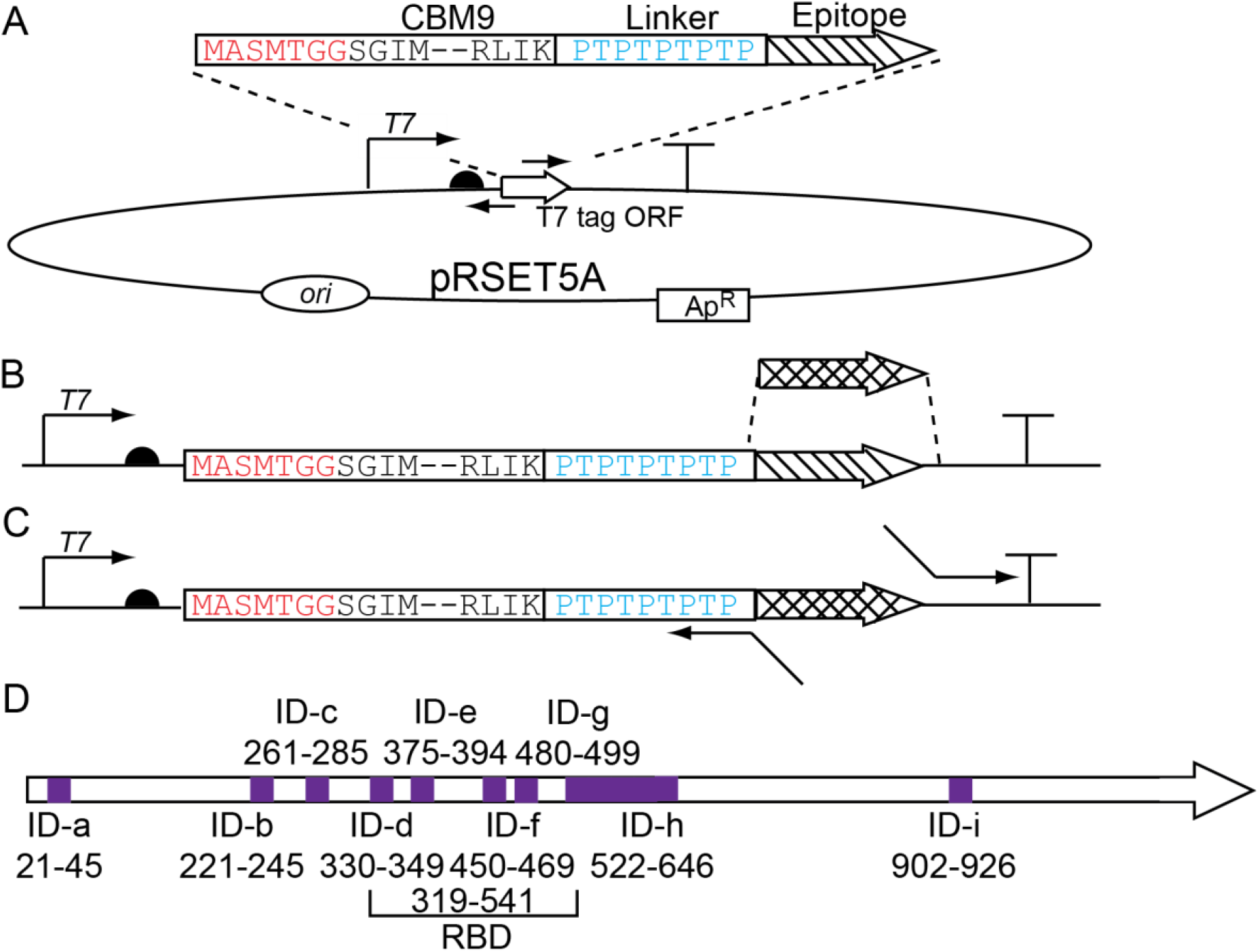
Strategies used to clone SARS-CoV-2 epitopes as fusions to CBM9. A. Initial clones were made by amplifying pRSET5A by inverse PCR, and ligating the plasmid amplicon to synthetic DNA encoding CBM9 with a linker (abbreviated to save space) fused to spike protein epitope ID-C, ID-F, or ID-H3. B. To create fusion clones of ID-A, B, D, E, G, H1 and I, synthetic DNA encoding just the epitope regions replaced the ID-C encoding region. C. To create the clones CBM9-(PT)_4_P, and N, primers with long overhang regions were used in an inverse PCR reaction using pRSET5A::*CBM9-id-a* to exchange the epitopes fused to CBM9. D. Representation of linear ID-A through ID-I regions with the amino acid numbers of the SARS-CoV-2 spike protein recognized by antibody from COVID-19 convalescent sera, as described by Zhang et al. (10); RBD is the receptor binding domain. In Panels A-C the bent arrow indicates T7 promoter region; half circle indicates T7 ribosome binding site; “T” symbol represents transcriptional terminator.

### Fusion protein isolation and analysis

Frozen aliquots of seed cultures were prepared and 0.4 mL of the stocks were added to 20 mL of modified auto-induction ZYM-5052 media that lacks an added carbon source (1% N-Z amine, 1% yeast extract, 25 mM Na_2_HPO_4_, 25 mM KH_2_PO_4_, 50 mM NH_4_Cl, 5 mM Na_2_SO_4_, 2 mM MgSO_4_, 100 μM FeCl_3_) (26); the media was supplemented with carbenicillin (250 μg/mL) and chloramphenicol (10 μg/mL). These cultures were grown at 24 °C for 16 h. The A_600_ was determined and the culture was added to 100 mL fresh broth with antibiotics in a 1 L flask to give an A_600_ of 2. IPTG was added to a final concentration of 2 mM, and the culture was incubated with shaking (250 RPM) for 3 h at 30 °C. The cultures were cooled on ice for 15 min, and then subjected to centrifugation for 10 min at 16,000 x *g*. The cell pellet from each 50 mL of culture was suspended in 5 mL of solution B (500 mM NaCl, 10 mM MgCl_2_, 0.5% CHAPS, 50 mM potassium phosphate, pH 7.0, 100 μg/mL lysozyme). 3 g of glass beads (≤106 μm, Sigma), and 5 μL Benzonase® nuclease (Sigma) were added and the cell suspension was vortexed vigorously for two 1 min intervals, and then subjected to centrifugation for 10 min at 16,0000 x *g*. The supernatant was added to 0.5 g of cellulose powder (Sigma, cat # 435236) that had previously been equilibrated with solution A (500 mM NaCl, 50 mM potassium phosphate, pH 7.0) in a 15 mL conical centrifuge tube. The tube was rocked for 1 h at room temperature, and the cellulose powder was pelleted by centrifugation for 2 min at 4,000 x *g*. The cellulose resin was washed three times by adding 12 mL of solution A, rocking for 15 min, and separation of the cellulose from the washing solution. This was followed by three washes using solution C (150 mM NaCl, 50 mM potassium phosphate, pH 7.0). Following the last wash with solution C, the cellulose was suspended in elution buffer (1 M glucose, 15 mM NaCl, 10 mM Tris, pH 7.6), and rocked for 15 min. Centrifugation was applied to pellet the cellulose, and the supernatant was removed and added to a Vivaspin® 6, 10 kDa MW cut off (Sartorius) protein concentrator to change the buffer to 150 mM NaCl, 10 mM Tris, pH 7.6.

For SDS-PAGE analysis of the CBM9 fusion protein-expressing cultures, 1 mL of cultures were centrifuged and the cell pellet suspended in 150 μL of Bug Buster (EMD Millipore), supplemented with lysozyme and Benzonase® nuclease as described above. Cell extracts were analyzed by SDS-PAGE using MOPS-SDS running buffer and TruPAGE™ gels (Sigma).

### Mass spectroscopy analysis of recombinant proteins

Liquid chromatography–mass spectrometry for the determination of intact molecular weight of isolated CBM9 fusion proteins was conducted by the University of Victoria/Genome BC Proteomics Centre. Details of the methods are presented in the Supplemental Materials.

### Commercial recombinant proteins and antibodies

The following recombinant SARS-CoV-2 proteins (18) that were expressed in *E. coli* were procured from the MRC Protein Phosphorylation and Ubiquitination Unit Reagents and Services at the University of Dundee (Dundee, Scotland; https://mrcppu-covid.bio/): MBP-Spike RBD amino acids (aa) 319-541 (DU 67753); GST-NSP1 aa 1-180 (DU 66413); GST-NSP2 aa 1-638 (DU 66414); NSP13 aa 1-601 (DU 66417); GST-NSP14 aa 1-527 (DU 66418); GST-NSP15 aa 1-527 (DU 66419); GST-Membrane Protein aa 1-222 (DU 67699); and GST-Nucleocapsid Protein aa 1-419 (DU 67726).

Rabbit polyclonal antibodies (Kinexus Bioinformatics) directed against synthetic peptides based on SARS-CoV-2 proteins were used in dot blots and are as follows: Spike aa 333-353 (NNCOV2S-5); Spike aa 450-467 (NNCOV2S-6); Spike aa 480-494 (NNCOV2S-7); Spike aa 505-524 (NNCOV2S-1); Spike aa 566-581 (NNCOV2S-9); Spike aa 574-588 (NNCOV2S-10); Membrane aa 3-23 (NNCOV2M-1); Nucleocapsid aa 156-170 (NNCOV2N-1); ORF1a aa 151-168 – NSP1 (NNCOV2-1A-1); ORF1a aa 735-750 – NSP2 (NNCOV2-1A-2); ORF1b aa 5606-5619 – NSP13 (NNCOV2-1B-1); ORF1b aa 6053-6071 – NSP14 (NNCOV-1B-2) and ORF1b aa 6713-6735 – NSP15 (NNCOV2-1B-3).

### Human Serum Samples

The process of the use of human blood products for the investigation of COVID-19 received an ethical approval by Veritas Independent Review Board Inc. (Saint-Laurent, QC, Canada), IRB Protocol 16567-09:39:354-06-2020. The majority of blood serum samples were generated from blood donations of volunteers. A small number of serum or plasma samples from positive tested donors and all pre-COVID-19 samples were acquired by purchase (Precision for Medicine, Norton, MA, USA; Innovative Research, Novi, MI, USA; AllCells, Alameda, CA, USA) or by donation (CureImmune, Vancouver, BC, Canada).

The blood samples were collected from persons that had been confirmed as positive using a PCR genetic test (designated as “COVID”), those that showed symptoms similar to COVID-19 but were not tested (designated as “sick”), those that were healthy and asymptomatic (designated as “Control”) and those that were healthy and donated prior to April 2019 (“pre-COVID”). For recovery of the serum, the collected blood was allowed to clot at room temperature. After about 2 h, possible clots sticking on the walls of the tubes were released by scraping with pipette tips around the inside walls of the collection tubes before the samples were centrifuged at 2000 x *g* for at least 15 min. When larger blood volumes (up to 10 mL) were processed, the top layer containing the serum was transferred into a syringe and sterile filtered into storage tubes. Sera from small blood samples (less than 500 μL) were transferred directly into the storage tubes without filtration. The recovered serum was stored at −80 °C.

All of the preparations of recombinant proteins were robotically spotted as 0.24 μL of a ~ 8 μM final concentration (except GST-NSP2 SARS CoV2 [DU 66414] in Spot B1 was printed at a 6.5 μM concentration) on to nitrocellulose membranes. The blots were washed three times with TBS (aqueous solution of 20 mM Tris-base and 250 mM NaCl; pH 7.5). These dot blot arrays were blocked with 2.5% BSA in T-TBS for 30 min. After washing the membranes twice with T-TBS (TBS with 0.05% Tween 20), the arrays were incubated with either affinity-purified rabbit polyclonal antibodies against SARS-CoV-2 protein sequences at 2 μg/mL or serum from recovered COVID-19 patients and healthy controls at 1:200 dilution in T-TBS. The incubation was carried out at 4 °C overnight. To detect the bound antibodies, the arrays were washed with T-TBS three times followed by the incubation with goat anti-human IgG+IgA+IgM pAb or donkey-anti-rabbit IgG (HRP conjugates, 1:20,000 dilution; Jackson ImmunoResearch, West Grove, Pennsylvania, USA). After 30 min incubation, the arrays were washed six times with T-TBS, once with 0.125 M NaCl and rinsed twice with water. The bound secondary antibody was visualized by enhanced chemiluminescence (ECL) on a Bio-Rad FluorS-Max scanner. The ECL scan was performed at a scanning time of 300 s. Eight images of the scanned array were generated during that scanning time.

### Accession numbers and availability of recombinant clones

The nucleotide sequences for all of the clones described in this work were deposited with GenBank (CBM9-(PT)_4_P, MZ322548; CBM9-ID-A, MZ322549; CBM9-ID-B, MZ322550; CBM9-ID-C, MZ322551; CBM9-ID-D, MZ322552; CBM9-ID-E, MZ322553; CBM9-ID-F, MZ322554; CBM9-ID-G, MZ322555; CBM9-ID-H1, MZ322556; CBM9-N, MZ322557). The recombinant clones expressing CBM9-(PT)_4_P, CBM9-ID-F, CBM9-ID-H1 and CBM9-N have been deposited with AddGene (https://www.addgene.org/).

## RESULTS AND DISCUSSION

### Expression of SARS-CoV-2 protein fragments as fusions to the CBM9 module

Pure proteins or fragments of proteins from the SARS-CoV-2 virus can be used to test for the presence of anti-viral antibodies in serological assays. The amount of antigen needed can be minimized if the fragments of the viral proteins that elicit an antibody response are known and these specific fragments are used in the serological assay. The goal of our research was to test if SARS-CoV-2 protein fragments known to elicit a human antibody response could be produced inexpensively using a universally available microbial expression system. The SARS-CoV-2 spike protein has been recombinantly produced in a number of hosts, including engineered HEK-293 cells, insect cells, and insect larvae (17, 27, 28). However, these expression systems require growth media and bioreactors that are orders of magnitude more expensive than the material and equipment used for microbial expression systems. Therefore, we felt that there was value in developing an inexpensive microbial expression system that can produce SARS-CoV-2 protein fragments in substantial amounts.

Our strategy was to use a standard *E. coli* T7 RNA polymerase approach to drive high levels of mRNA transcription, and use a proven carrier protein module, a thermophilic family 9 carbohydrate-binding module (CBM9), to carry the SARS-CoV-2 fragment at the C-terminus of the fusion protein (Fig. 1 A-C). Kavoosi et al. (21) showed that the CBM9 module expresses at high levels, even when a protein was fused to the C-terminus. In a separate work (22), these same researchers showed that linking CBM9 to a protein with a proline-threonine rich linker ([PT]_4_P-IEGR) resulted in a fusion protein that was resistant to protease attack by endogenous *E. coli* proteases (22). Thus, we adopted the use of (PT)_4_P as a linker between the CBM9 and SARS-CoV-2 spike protein fragments, and an example of the gene organization is shown in Supplemental Fig. S1.

The bulk of the studies on the antibody response to SARS-CoV-2 have been conducted using overlapping synthetic peptides corresponding to SARS-CoV-2 proteins, primarily the spike protein, but sometimes several proteins or the entire proteome. At the time that we initiated our studies, the work of Zhang et al. (10) was one of the more comprehensive analyses of the human antibody response in COVID-19. These researchers identified nine regions in the spike protein, designated ID-A through ID-I (Fig. 1D), which were recognized by antibodies from COVID-19 convalescent sera, and we chose these regions as well as one nucleocapsid protein epitope to clone and express. An SDS-PAGE analysis of the expressed CBM9-spike protein fusions (Supplemental Fig. S2) indicated which fusion proteins were most resistant to degradation by *E. coli* and we chose a sub-group of the clones to study further (abandoning ID-H, ID-H2, ID-H3 and ID-I). Table 1 lists the amino acid sequences of the encoded spike (and one nucleocapsid [“N”]) protein regions in the clones we constructed and chose to study further, and the specific epitope regions of the spike protein identified by Zhang and co-workers (10) are underlined.

**Table 1.**
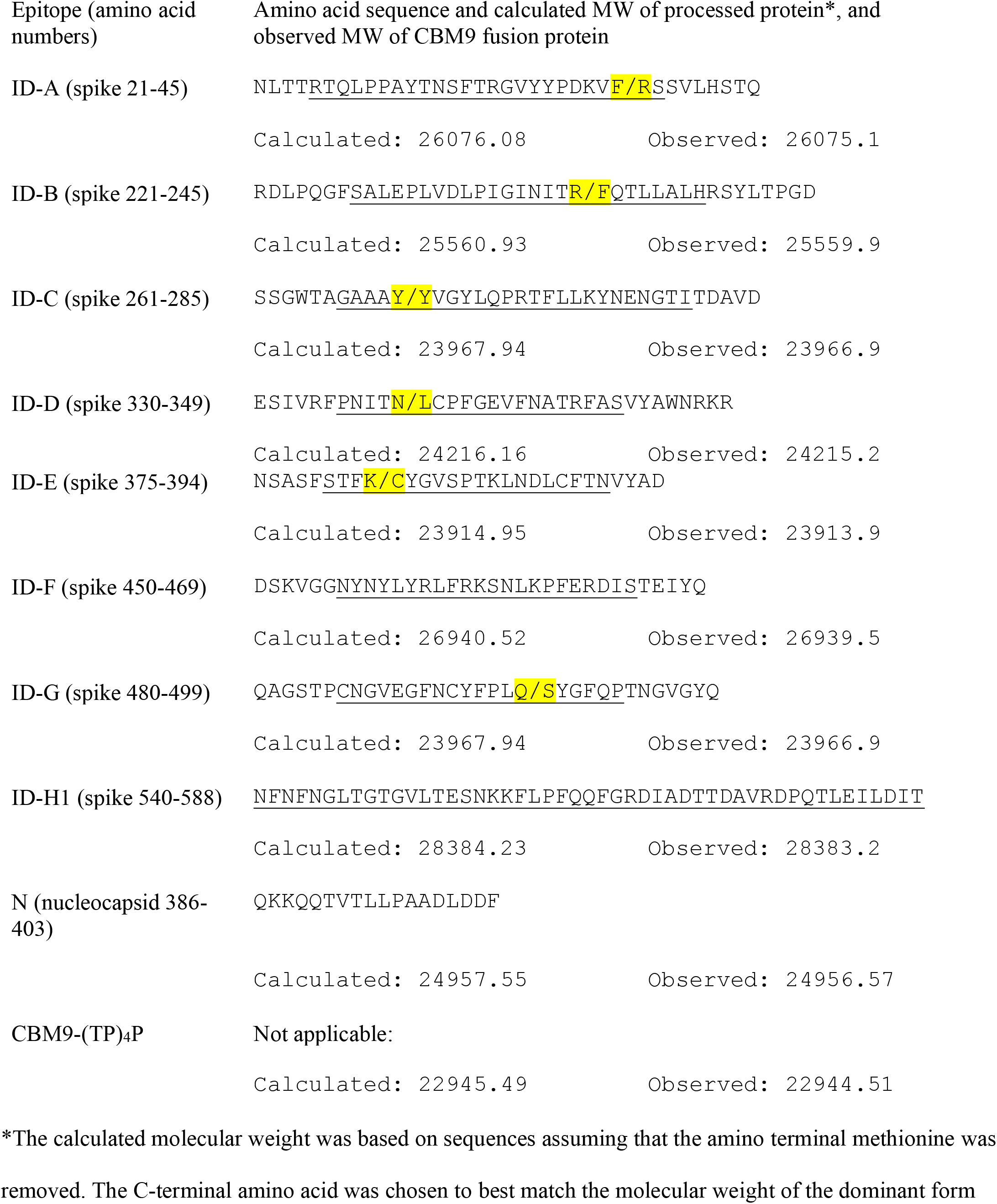
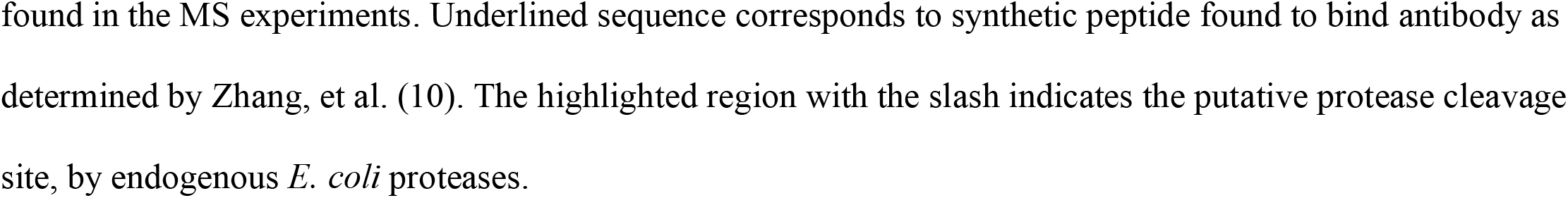
Amino acid sequence of B-cell epitopes cloned as fusions to CBM9, and location of putative protease cleavage sites.

### Purification and mass spectroscopy analysis of CBM9 fusion proteins

From the recombinant CBM9 fusion clones that we chose, we expressed and isolated the recombinant protein using powdered cellulose in a batch purification. The resulting purified proteins were subjected to mass spectroscopy analysis to determine the molecular weight of the dominant purified product. The results (Table 1) indicated, as expected, that all products had the N-terminal methionine removed. Most cloned products were processed, presumably by endogenous *E. coli* proteases, so that some portion of the C-terminal end was removed, effectively removing a few to several amino acids of the spike protein fragment. However, clones expressing CBM9-(PT)_4_P, CBM9-ID-F, CBM9-H1 and CBM9-N produced, as the dominant purified product, proteins that were 1 Dalton smaller than the predicted monoisotopic product. Since the CBM9-(PT)_4_P dominant product was 1 Dalton smaller than predicted, we interpreted this to mean that an unidentified chemical modification occurs, many of which are documented (29), on the CBM9 module to remove one atomic mass unit. The clone expressing CBM9-ID-A was processed so as to remove only two amino acids from the B-cell epitope identified by Zhang et al. (10).

For further work we chose *E. coli* clones that expressed intact fusion proteins of CBM9 and fragments of SARS-CoV-2 proteins as the dominant recombinant products. The expression of clones CBM9-(PT)_4_P, CBM9-ID-H1 and CBM9-N are shown in Fig. 2A and the purified products in Fig. 2B; for comparison, the carrier protein module CBM9 is also shown. By comparing the staining intensity of the protein band in the cell extracts to the band of purified CBM9-N, we estimated that the clones expressed recombinant product at levels of about 100 mg/L upon IPTG-induction, and this is consistent with the 200 mg/L estimates of Kavoosi et al. (21). These experiments were performed using standard research growth flasks at an A_600_ of less than 10, and presumably the levels of recombinant protein that are produced could be significantly increased using an optimized fed-batch bioreactor protocol.

**Figure 2.**
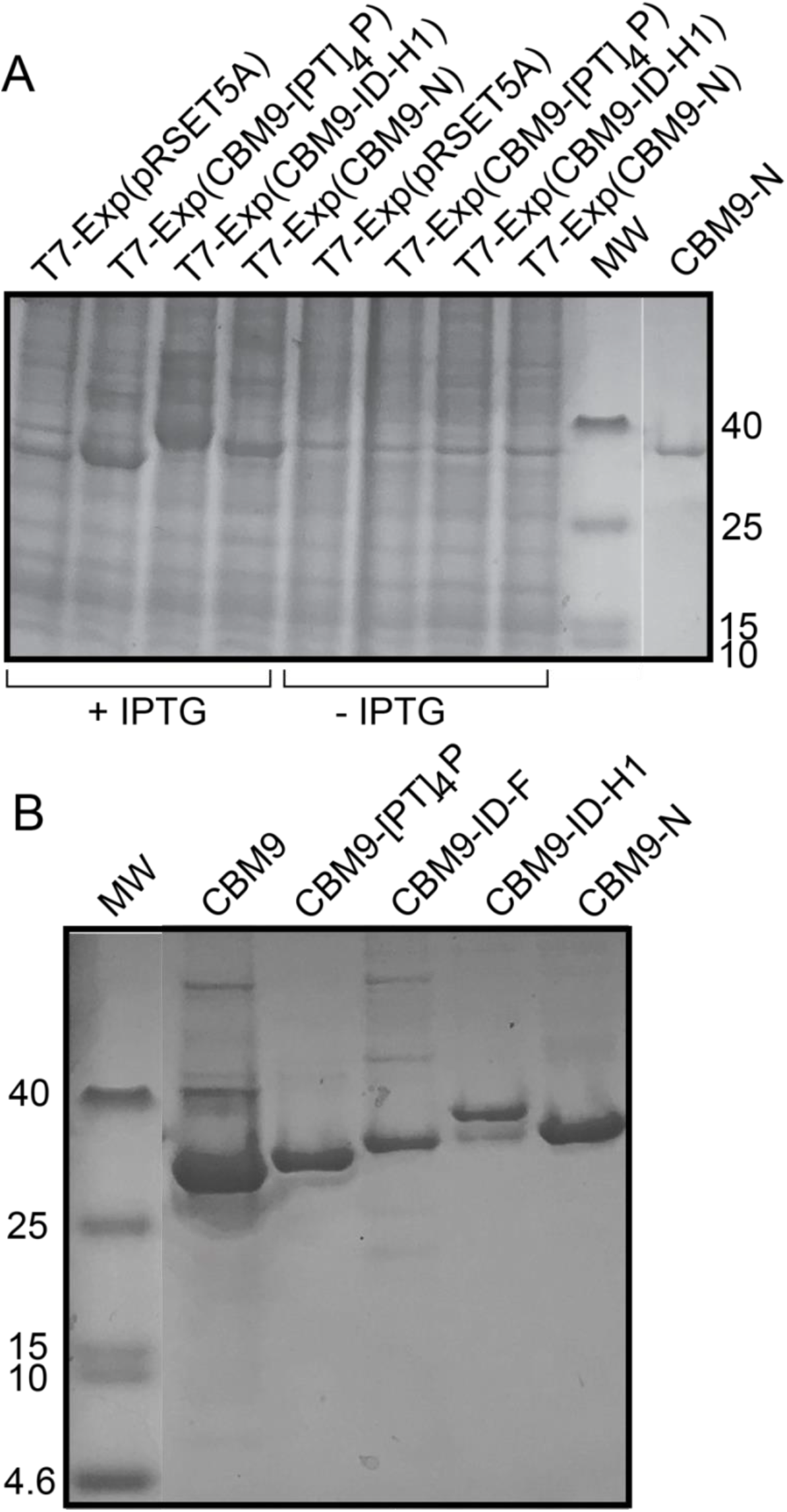
A. Cell extracts, equivalent to 33 μL of detergent-soluble material, of recombinants encoding CBM9 fusion proteins with and without IPTG induction. For comparison 1.3 μg of purified CBM9-N is shown in the last lane on the right. B. Samples of CBM9 fusion proteins purified by batch absorption to cellulose powder, after storage at 4 °C for a minimum of two weeks.

The recombinant fusion proteins were isolated by binding to cellulose powder, using batch purification (Fig. 2B). Following this, the CBM9 and the CBM9-ID-F samples were heated to 70 °C for 10 minutes and the CBM9-(PT)_4_P, CBM9-ID-H1 and CBM9-N samples were filtered sterilized, all before storing at 4 °C for at least two weeks (Fig. 2B). As well, samples were stored at −20 °C in 50% glycerol (Fig. 2A last lane on right, CBM9-N). All storage conditions preserved the integrity of the sample. However, heating to 70 °C seemed to generate small amounts of multimers of the protein, consistent with previously reported observations for hyperthermophilic enzymes (30).

We constructed a number of clones of the ID-H region (see Fig. S2C), and it was fortuitous that the CBM9-ID-H1 clone highly expressed a product that was largely resistant to *E. coli* proteases. The ID-H region (residues 522-646) partially overlaps with the RBD (residues 319-541) of the SARS-CoV-2 spike protein. In the 3D-structure oriented with the RBDs at the top, the ID-H1 region (residues 540-588) slightly overlaps with and lies below the RBD (Fig. 3A). It is possible that the CBM9-H1 recombinant product is resistant to proteases, while shorter than other CBM9 fusions that are susceptible, because the ID-H1 clone encodes a potential self-folding protein domain (Fig. 3B). The region encompassed by CBM9-ID-H1 includes amino acid sequences identified by several groups as B-cell epitopes, as defined by synthetic peptides that are recognized by convalescent sera from COVID-19 patients (Fig. 3C).

**Figure 3.**
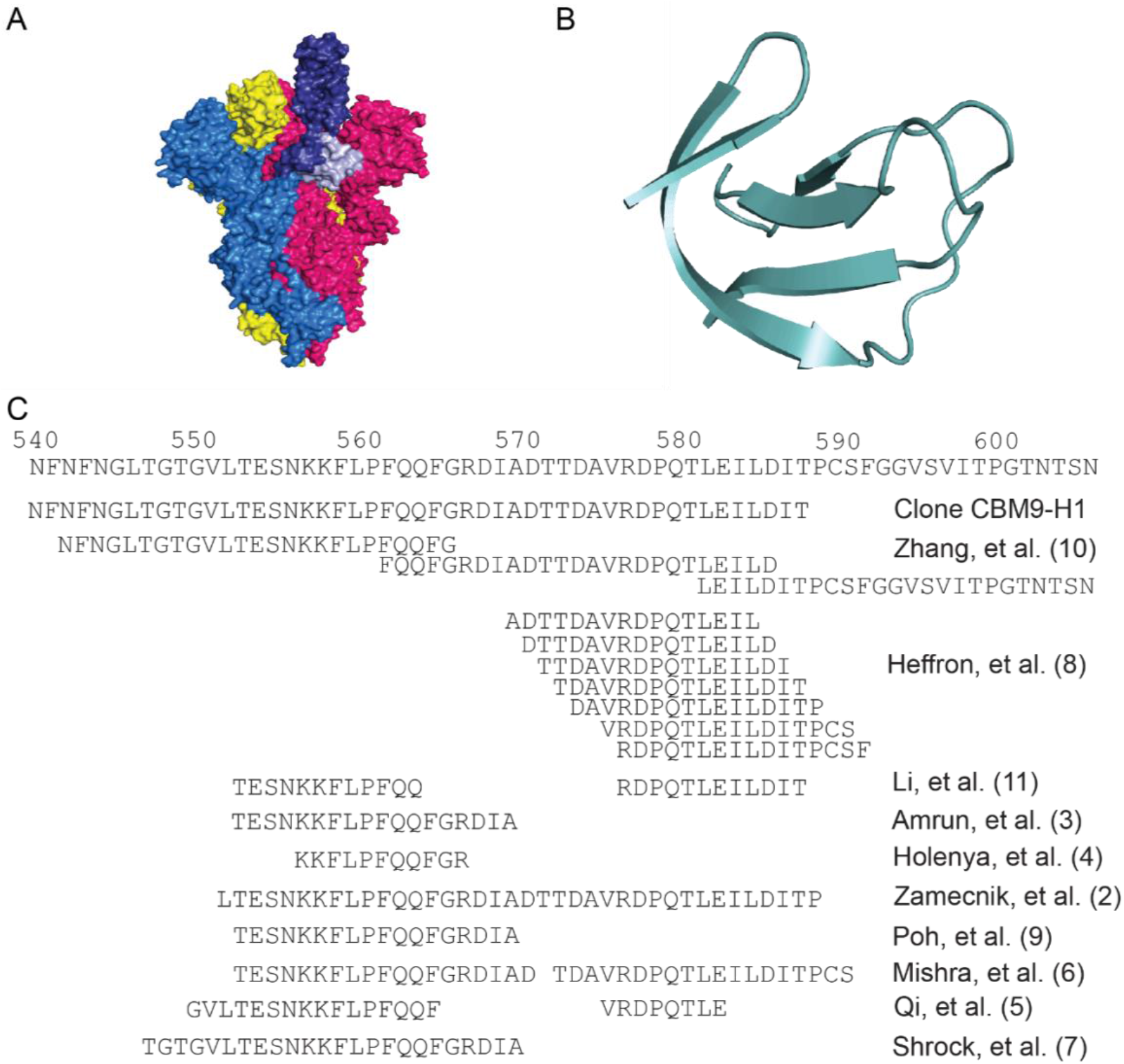
A. Surface topology view of an electron microscopy structure of the Spike protein trimer in the open state (PBD: 6VYB) (33), created with PyMOL (34). The S protein subunit in the up position is colored in blue and the subunits in the down position are colored in yellow and pink. The RBD of the blue subunit is shown in dark blue and the H1 region in light blue, though it is important to note that some residues of the RBD are missing in the electron microscopy structure. B. Cartoon representation of the H1 region of the S protein (PDB: 6VYB) (33), created with PyMOL (34). C. Immunodominant region of SARS-CoV-2 spike protein. Several groups have identified the region that encompasses approximately amino acids 540-600 as a region that elicits antibody response following infection.

Interestingly, the region encompassed by the H1 clone does not encompass any of the amino acid changes found among any of the variants of concern (12), but it does contain a putative protective epitope. Poh et al. (9) found that titer of antibody in sera from COVID-19 convalescent patients that reacted with a peptide corresponding to amino acids 562-579 of the spike protein correlated with the amount of *in vitro* pseudovirus neutralization. Further, when the neutralizing sera was depleted of reactivity against the peptide representing amino acids 562-579 the neutralization activity fell sharply. Such evidence indicates that a strategy to elicit antibodies against this region may be an effective way of protecting against variants with amino acid changes in the RBD of the SARS-CoV-2 spike protein.

### CBM9-SARS-CoV-2 fusion proteins react with rabbit anti-spike protein sera and human sera

As stated above, we used proline-threonine flexible linker regions to join the CBM9 module to SARS-CoV-2 spike protein and nucleocapsid protein regions. In this research the purpose of the linker was to allow the SARS-CoV-2 protein fragment to be accessible to antibody binding. To determine if the linker accomplished this, we reacted purified CBM9-(PT)_4_P, CBM9-ID-F, CBM9-ID-H1 and CBM9-N with purified rabbit antibodies that had been raised to different portions of SARS-CoV-2 proteins (Fig. 4) and human sera (Fig. 5 and Fig. 6). In a semi-quantitative dot blot assay, we found that rabbit antibodies raised against the appropriate fragments of the SARS-CoV-2 spike protein reacted strongly with CBM9-H1 (Fig. 4F, 4G), but only weakly, or not at all, with antibodies directed against other regions of the spike protein. Reaction of the appropriate antibodies with CBM9-ID-F was moderately strong (Fig. 4C), and likewise poor or not detectable with the other antibodies. We had access to a small sampling of sera from COVID-19 confirmed (n=7), COVID-19 suspected (n=13), and healthy individuals (n=20) (Fig. 5 and Fig. 6). While this small sample set size and the dot blot assay cannot provide an epidemiological story, the results did show that human sera clearly reacted with the CBM9 fusions carrying ID-F and the nucleocapsid epitope. Many sera samples from both sick and healthy individuals reacted strongly with ID-F, indicating that this region may be similar in other coronaviruses or may be similar to another commonly encountered antigen. Surprisingly, the sera from several individuals, both healthy and ill, reacted apparently more strongly with CBM9-ID-F (spike amino acids 450-469) than with the MBP-RBD fusion protein (spike amino acids 319-541), even though the latter encompasses the ID-F region. These results may reflect a property of the antigen, such as accessibility of the SARS-CoV-2 portion to the antibody; or it may be that the ID-F region is an especially immunogenic region of the RBD. Overall, with this small sample of sera there was little difference between the patterns of reactivity of the sera from the sick and healthy groups. This high detection of anti-spike and anti-nucleocapsid immunoreactivity in serum samples from healthy individuals is consistent with previous studies using two different serological tests developed by Mesoscale Devices and Kinexus (31).

**Figure 4.**
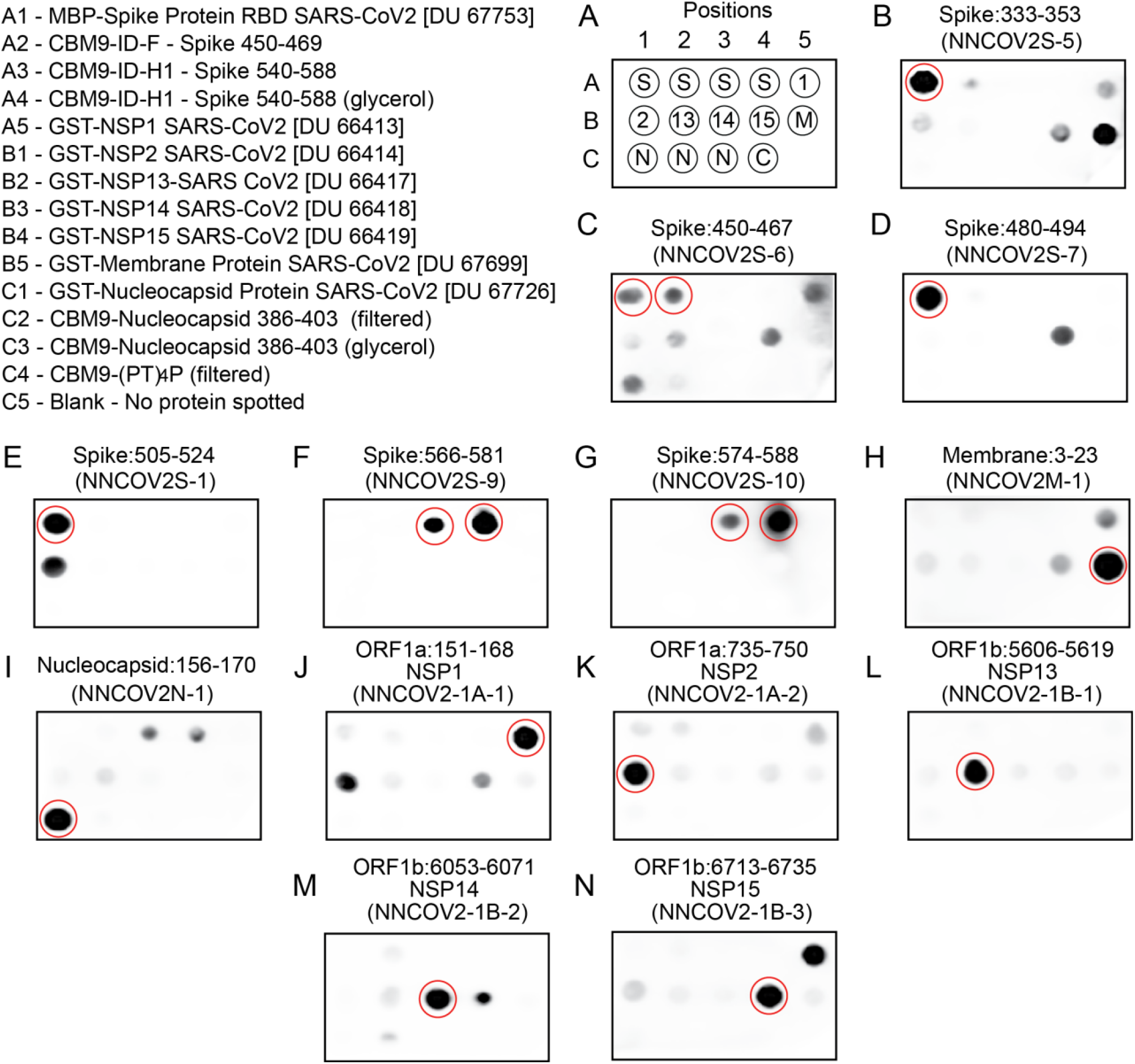
Dot blotting analysis of SARS-CoV-2 recombinant proteins with rabbit polyclonal antibodies for diverse SARS-CoV-2 proteins. Expected target positions of SARS-CoV-2 proteins for each antibody are circled. Identification and location of each recombinant protein is shown in Panel A.

**Figure 5.**
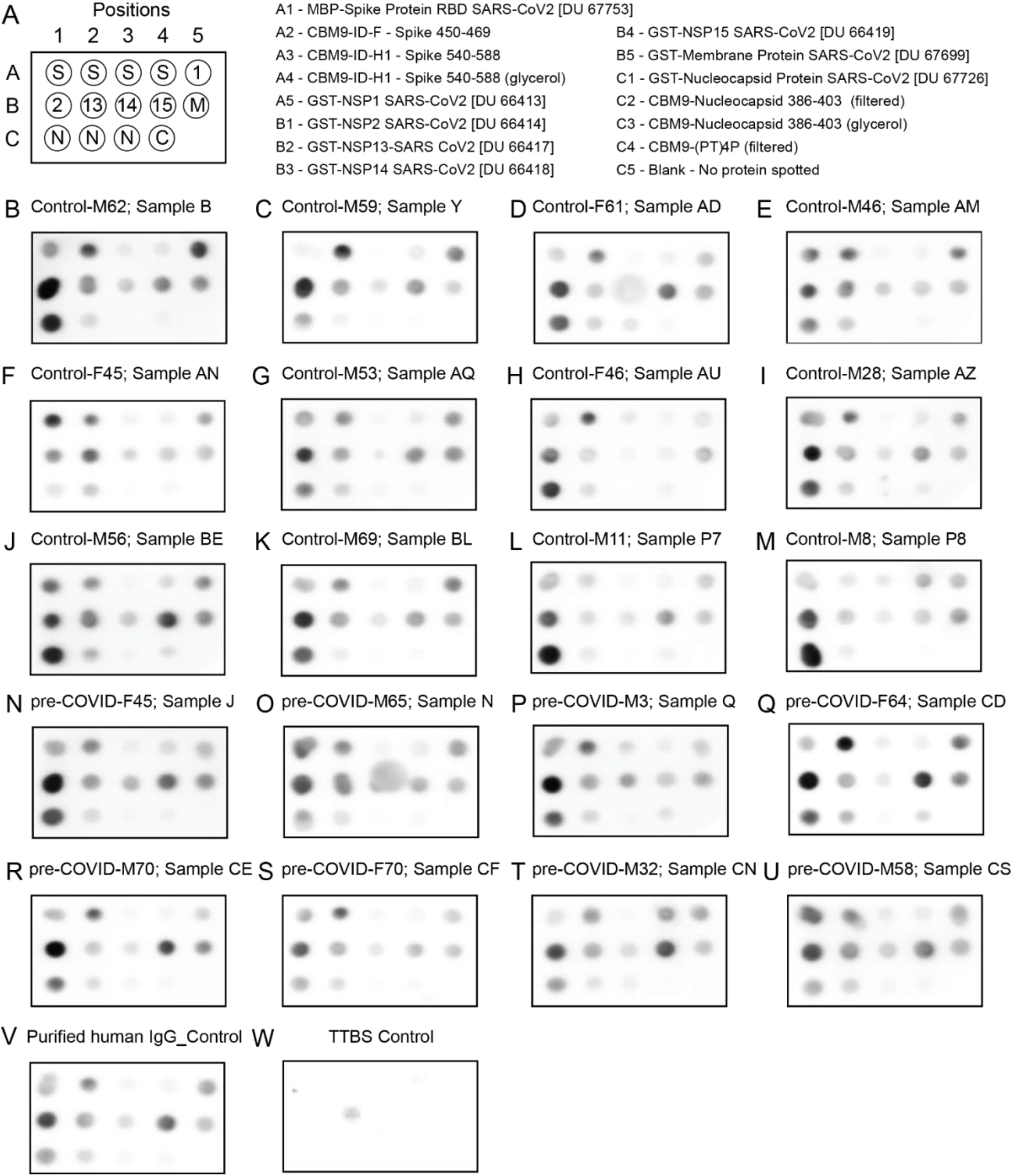
Dot blot of proteins with sera from pre-COVID and healthy controls. Panels B-M, “Control” samples correspond to healthy individuals whose serum samples were collected in 2020. Panels N-U, “pre-COVID” samples were from healthy individuals whose serum samples were retrieved prior to April 2019. Individuals are also identified by sex (M for male and F for female) followed by age in years. The MBP-spike RBD protein (spot position A1) includes amino acids 319-541 of the spike protein.

**Figure 6.**
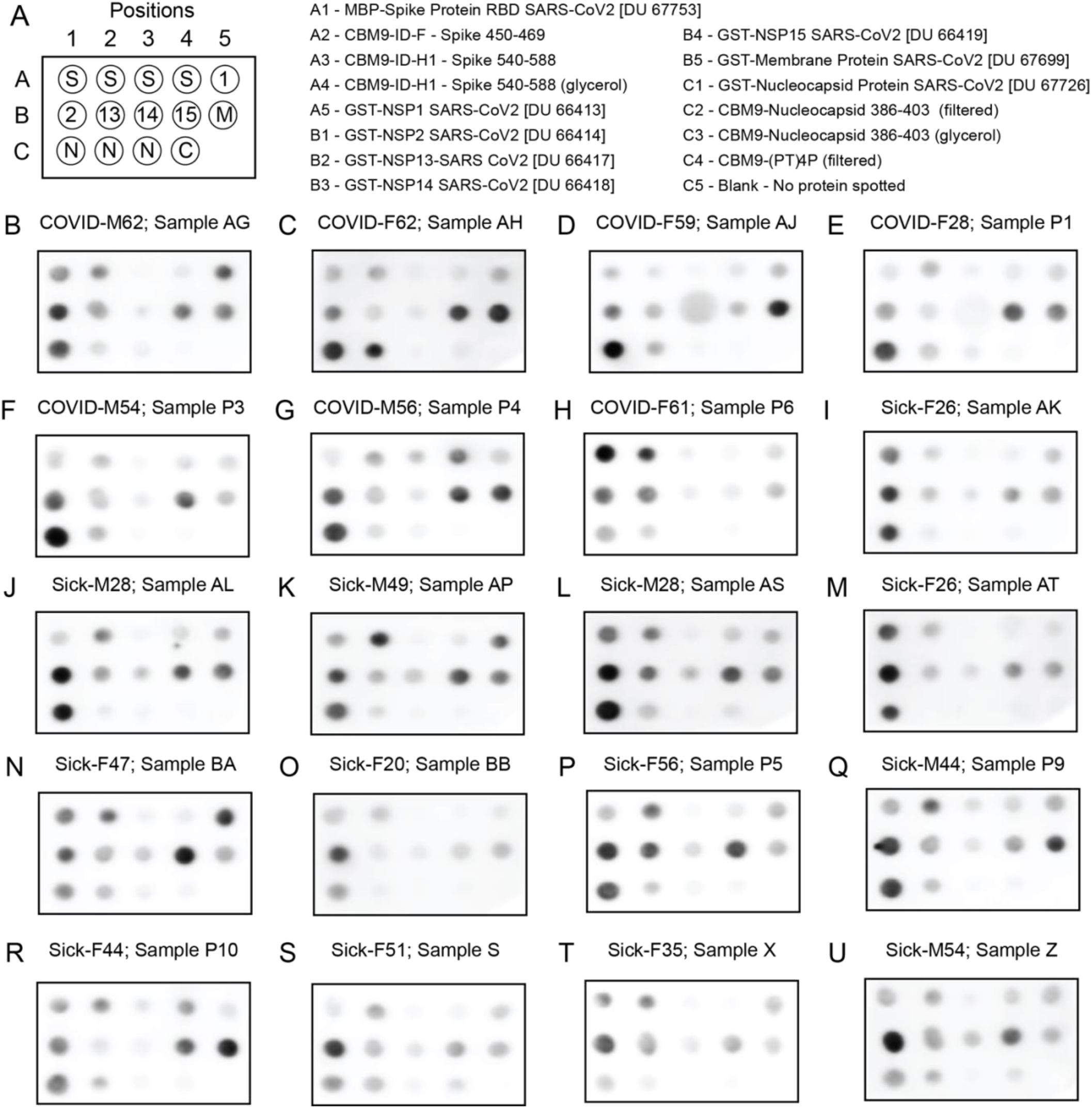
Dot blot of proteins with sera from COVID-19 and sick individuals. Panels B-H, “COVID” samples correspond to individuals that PCR-tested positive for SARS-CoV-2 RNA and whose serum samples were collected in 2020. Panels I-U, “Sick” samples were from individuals who had COVID-19 symptoms and whose serum samples were collected in 2020. Individuals are also identified by sex (M for male and F for female) followed by age in years. The MBP-spike RBD protein (spot position A1) includes amino acids 319-541 of the spike protein.

In this work, we have demonstrated that fragments of the SARS-CoV-2 virus, fused to the CBM9 module through a flexible linker, could be produced at high levels - about 0.1 g/L - using universally available equipment with inexpensive materials. The costs of cultivating *E. coli* are 10-to 100-fold less expensive than the costs of growing non-microbial eukaryotic cells, which are the usual hosts for expressing SARS-CoV-2 antigens. Further, the cost of using cellulose powder for affinity purification of CBM9 fusion proteins is about 100-to 1,000-fold less than using the conventional immobilized nickel resin or a combination of traditional protein purification columns, such as ion-exchange with size exclusion resins. Lastly, while we described the production and isolation of CBM9 fusion proteins, there may be applications that require the separation of the CBM9 module from the SARS-CoV-2 fragment. Often the components of a fusion protein are separated using a highly specific protease cleavage site. Indeed, using this approach Kavoosi et al. (21) found that a CBM9-GFP fusion protein remained largely intact even in the absence of protease inhibitors, unless cleaved with factor Xa whose cleavage site was incorporated into the linker. While specific proteases may be found that function with a specific CBM9 fusion, with or without addition of the cleavage site into the linker, it is also possible to use chemical treatments that cleave with an acceptable level of specificity. For example, in the case of recombinant product CBM9-ID-H1, 45 amino acids of the 49 amino acid SARS-CoV-2 fragment could be released by treatment with hydroxylamine which attacks asparaginyl-glycine bonds (32) as CBM9 lacks an arginine-glycine sequence. Thus, the use of CBM9-SARS-CoV-2 protein fragment fusions allows for the economical production of antigens to be used for a variety of purposes, including in COVID-19 serological assays.

## ACKNOWLEDGEMENTS

We thank Jeff Wang for aid in the drawing of blood samples from the study participants. Our thanks go also to Akshra Atrey for the probing of the dot blot arrays.

